# High-affinity PQQ import is widespread in Gram-negative bacteria

**DOI:** 10.1101/2024.06.04.597491

**Authors:** Fabian Munder, Marcos Voutsinos, Klaus Hantke, Hari Venugopal, Rhys Grinter

## Abstract

PQQ is a soluble redox cofactor used by diverse bacteria to oxidise fuel compounds as a source of electrons for the respiratory chain. Many Gram-negative bacteria that encode PQQ-dependent enzymes do not possess the biosynthetic machinery for its production and instead obtain it from the environment. To achieve this the bacterium *Escherichia coli* uses the TonB-dependent transporter PqqU as a high-affinity PQQ importer, allowing it to use PQQ at an external concentration as low as 1 nM. Here, we show that PqqU achieves this by binding PQQ with a very high affinity. Using cryo-electron microscopy we determine the structure of the PqqU-PQQ complex at a resolution of 1.99 Å, revealing that the extracellular loops of PqqU undergo significant conformational changes upon PQQ binding, which captures the cofactor in an internal cavity. This cavity likely facilitates an airlock-style gating mechanism that prevents non-specific import through PqqU. Using structural modelling we show that the change in PqqU structure upon PQQ binding precludes the binding of bacteriophage, which targets it as a cell surface receptor. Guided by the PqqU-PQQ complex structure we use phenotypic analysis to identify the amino acids essential for PQQ import and leverage this information to map the presence of PqqU across Gram-negative bacteria. This reveals that PqqU is encoded by Gram-negative bacteria from at least 22 phyla from diverse habitats, including those found in aquatic, soil, host-associated, and extreme environments. This indicates that PQQ is a ubiquitous nutrient in many environments, and an important cofactor for bacteria that adopt diverse lifestyles and metabolic strategies.

**Significance Statement:** Many enzymes form complexes with molecules called cofactors to perform their function. PQQ is a cofactor used by bacterial enzymes that provide energy by breaking down food molecules. While some bacteria make their own PQQ, other bacteria use the transport protein PqqU to bind PQQ from the environment and import it into their cells. We show that PqqU binds PQQ very tightly, allowing bacteria to acquire it at very low concentrations. Using cryo-electron microscopy we image the PqqU-PQQ complex on an atomic level, revealing how PQQ is bound so tightly. Using this the information to analyse microbial genomes, we show that PQQ scavenging is employed by diverse bacteria, implying that PQQ is an important common good of diverse microbiomes.

## INTRODUCTION

Pyrroloquinoline quinone (PQQ) is a soluble quinone redox cofactor used by a diverse family of calcium- and lanthanide-dependent dehydrogenases referred to as quinoproteins ^1-3^. Quinoproteins have been shown to oxidize substrates including sugars, alcohols, and aldehydes, and given the diversity of the family is likely that other substrate specificities exist ^4-9^. Quinoproteins are important metabolic enzymes in many environments, with PQQ-dependent methanol dehydrogenases the most abundant proteins found in some soil proteomes ^10^. These enzymes are predominantly found in Gram-negative bacteria, where they are localized to the periplasm and are sometimes membrane-associated ^1^. The periplasmic localization of quinoproteins allows for the oxidation of catabolic substrates without their import into the cytoplasm, with the resulting electrons delivered to the respiratory chain via the reduction of C-type cytochromes, membrane-bound respiratory quinones, or azurin ^1,9,11,12^.

Endogenous PQQ is synthesized in the cytoplasm before it is transported to the periplasm where it is bound and utilized by quinoproteins ^13-15^. However, the molecular mass of PQQ (330 Da) is below the diffusion limit of the Gram-negative outer membrane, and the unbound cofactor can be lost from the cell via diffusion ^16,17^. As a result, many PQQ-producing bacteria secrete PQQ under conditions where its production is induced ^18,19^. This secretion makes PQQ a common good of microbial communities, a fact which is exploited by bacteria like *Escherichia coli*, which utilize PQQ-producing enzymes but lack the biosynthetic enzymes for PQQ production ^20^.

The biosynthetic pathway for PQQ is complex, involving the production of the precursor peptide (PqqA), which is sequentially modified by four additional enzymes and a chaperone (PqqB-F) to generate the cofactor ^13^. Considering the complex and energy-intensive nature of PQQ biosynthesis, the loss of PQQ from the cell by diffusion is energetically unfavourable, and as such, producing bacteria employ systems to prevent the loss of PQQ from the cell, or to recapture PQQ from the environment ^16,17^. An example of this is the periplasmic PQQ binding protein PqqT recently identified in the Alphaproteobacterium *Methylobacterium extorquens* AM1. PqqT binds PQQ with a high affinity (Kd = 50 nM), helping to prevent the loss of free PQQ from the cell and facilitate its capture from the environment ^17^.

PQQ can cross the outer membrane and enter the bacterial periplasm by diffusion. However, this process is inefficient, and unless the external concentration of the cofactor is high it is unlikely to satisfy cellular requirements for the cofactor. Consistent with this, it was recently demonstrated that in *E. coli*, the TonB-dependent transporter (TBDT) PqqU (formerly YncD) enables the utilization of extracellular PQQ at concentrations as low as 1 nM, allowing phosphotransferase system (PTS) deficient cells to grow using the PQQ-dependent enzyme Gcd to oxidise glucose as the sole carbon/energy source ^16,21^. PqqU was not required for the growth of these cells at higher PQQ concentrations (100 to 3000 nM), indicating that the transporter is utilized for capturing the cofactor from the environment when concentrations are low. This provides strong evidence that PqqU is a high-affinity transporter of PQQ across the Gram-negative outer membrane ^16^.

In this current study, we show that consistent with its requirement for PqqU utilization at low concentrations, PqqU binds PQQ with very high affinity (Kd < 1 nM). To establish the structural basis for this high-affinity we determine the high-resolution CryoEM structure of the PqqU-PQQ complex at 1.99 Å. This structure shows that PqqU binds PQQ in a positively charged binding pocket, triggering significant conformational changes in the extracellular loops of PqqU preventing PQQ escape and blocking the binding of a PqqU targeting bacteriophage. We leverage this structural information to create PqqU binding site mutants and perform functional analysis to identify key residues for PQQ binding. We then utilize these key residues as a molecular signature to map the presence of PqqU across Gram-negative bacteria. We find that the capacity for high-affinity PQQ import via PqqU is present in at least 22 phyla from diverse environments, and it is widespread in Gammaproteobacteria, Bacteroidota, and Gemmatimonadetes. Only 23% of bacteria encoding PqqU also encode the biosynthetic machinery for PQQ biosynthesis, indicating that PQQ scavenging is an efficient strategy for diverse bacteria. Interestingly, PqqU producers often, but not always, encode putative quinoproteins, indicating that there may be additional uses for the co-factor outside this enzyme family. The high frequency of PqqU in diverse Gram-negative bacteria indicates that PQQ is an important nutrient for bacteria adopting diverse metabolic strategies and high-affinity PQQ import is a beneficial trait in many environments.

## RESULTS AND DISCUSSION

### PqqU binds PQQ with very high affinity

Previous work showed that PqqU is required for high-affinity uptake of the redox cofactor PQQ (Figure 1a) ^16^. However, this study did not provide direct evidence for interaction between this TonB-dependent transporter and PQQ, which would be required for import. To verify this interaction is occurring and to gain insight into its affinity and binding kinetics, we performed isothermal titration calorimetry (ITC), with PQQ injected into purified PqqU in the detergent LMNG in a micelle-depleted buffer (Figure S1). The ITC thermogram from this experiment shows negative saturable heats of binding indicative of an exothermic interaction between PqqU and PQQ (Figure 1b), with minimal heats observed for the buffer into PqqU control (Figure S2). Analysis of these data revealed a molar binding ratio of 0.74 PQQ molecules per PqqU, which likely indicates a 1:1 stoichiometry for this interaction, with not all PqqU modules in a binding competent state. A Gibbs free energy (ΔG) of -16.4 kcal/mol, was observed for PqqU-PQQ binding, which is largely driven by a favourable enthalpy of binding of -14.4 kcal/mol. This is consistent with the highly negatively charged nature of PQQ and the positively charged nature of the putative binding site of PqqU ^1,21^. A binding curve could not be accurately fitted to the integrated heats from the ITC titration, due to a lack of data across the inflection point, indicating that the interaction between PqqU and PQQ is of very high affinity, likely with a Kd of <1 nM (Figure 1c, Table 1). This is consistent with the biological role of PqqU as a high-affinity PQQ transporter ^16^.

**Figure 1:**
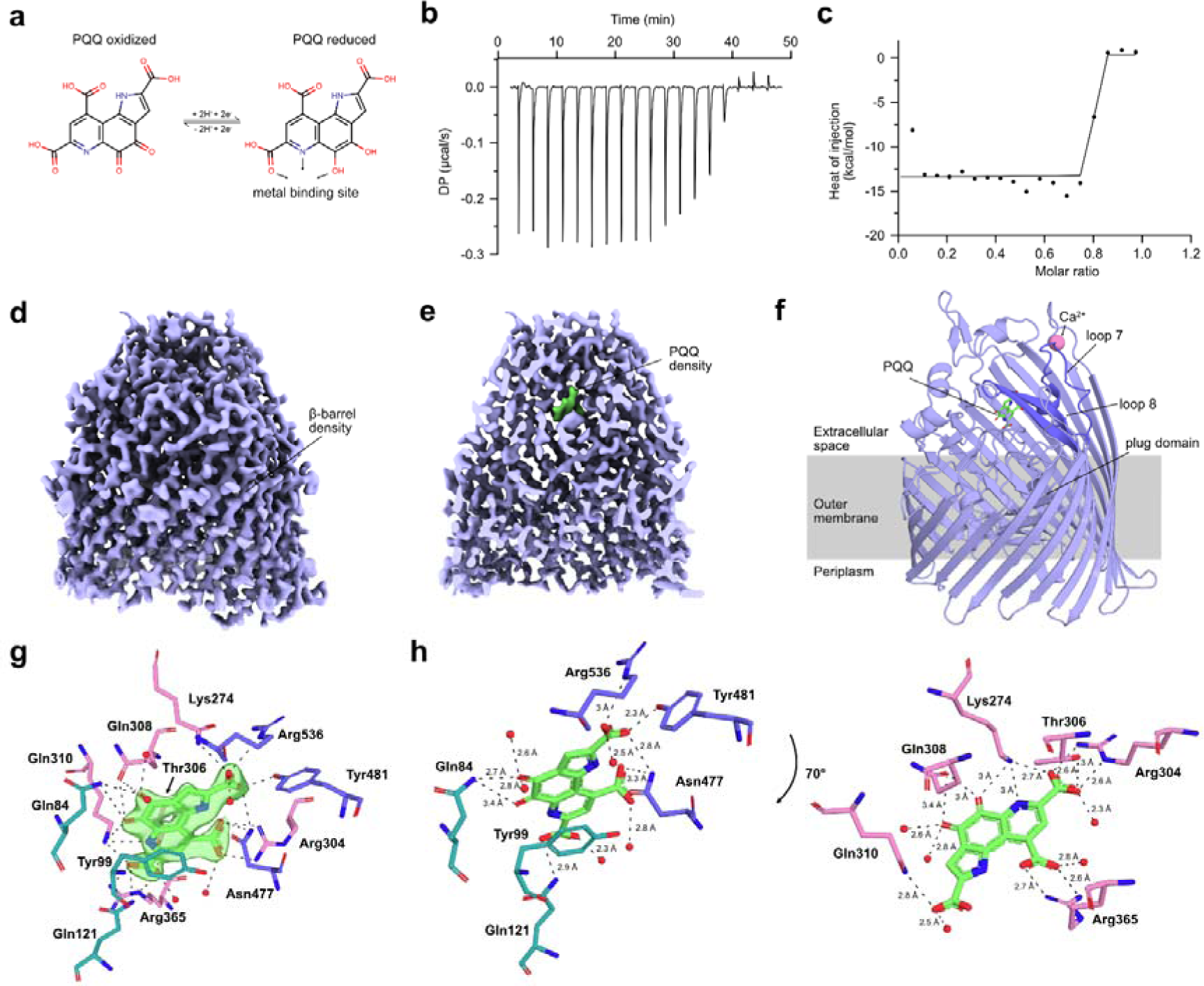
PqqU binds PQQ with very high affinity by enclosing it in an internal binding cavity. **a**, pyrroloquinoline quinone (PQQ) molecule visualized via a 2D stick model. PQQ changes from the oxidized form to the reduced form upon the two-electron transfer to both hydroxyl groups. The first aromatic ring contains nitrogen that together with two carbonyl groups coordinates a metal ion (Ca^2+^, or lanthanide) in PQQ-dependent enzymes (quinoproteins) **b**, a representative thermogram of 100 µM PQQ titrated into 20 µM PqqU over ∼50 minutes in 19 injections of 2 µl. The first injection (not included in graph) is 0.4 µl. **c**, negative peaks of injections integrated and plotted against the PqqU-PQQ molar ratio, excluding the first injection. Experiments were conducted in triplicate on a Malvern MicroCal PEAQ-ITC and analysed with the accompanying software. Baseline corrected data were plotted with GraphPad Prism. **d**, the PqqU Coulomb potential map reveals distinct structures of PqqU such as the characteristic β-barrel made of 22 individual β-strands. **e**, vertical cut of the PqqU Coulomb potential map to reveal density of PQQ. **f**, cryo-EM structure of PqqU with bound PQQ (green) in the closed conformational state at a global resolution of 1.99 Å. The extracellular loops 7 and 8 of PqqU undergo a conformational change which encloses the substrate binding pocket (dark blue). **g**, PQQ (green) with map density surrounded by the 12 active site residues coloured by location within PqqU (β-barrel = pink, loop 7 + 8 = blue, plug domain = green). Interaction distances between PQQ and residues/waters visualized by dotted lines. **h**, rotational view of PQQ with the 11 active site residues and bond lengths. Split into two parts for visual clarity based on location within the binding pocket.

### The high-resolution Cryo-EM structure of the PqqU-PQQ complex

To determine the structural basis for the high-affinity binding of PQQ by PqqU, we utilised cryo-electron microscopy (Cryo-EM) to resolve the structure of the PqqU-PQQ complex. PqqU in LMNG in micelle-depleted buffer was mixed with a twenty-fold excess of PQQ, before grid preparation and imaging on a Titan Krios microscope. Image processing and 3D reconstruction yielded high-quality maps of PqqU with a nominal resolution of 1.99 Å (Figure 1d,e, Figure S3 Table S2). These maps allowed us to model close to the complete polypeptide chain of PqqU from amino acid 39-699 out of a total of 700 amino acids, confirming that PqqU forms a monomeric 22-stranded β-barrel, with a transmembrane region enclosed in an LMNG micelle (Figure 1f, Figure S4, Movie S1). This high-resolution structure of PqqU (RMSD=0.957, 4030/5147 atoms), while globally similar to the PQQ-free structure previously solved by X-ray crystallography (YncD; PDB ID = 6V41), reveals significant conformational changes of the extracellular loops 8 and 7 and a general compaction of the transporter (Figure 2a, Movie S3). We observed unambiguous map density corresponding to a single PQQ molecule within the extracellular region of PqqU that was previously identified as a potential substrate binding pocket (Figure 1e)^21^. Interestingly, no density is observed for a metal ion in a complex with PQQ, indicating that PqqU does not co-transport the compound with a metal cofactor. However, the PqqU extracellular loop 1 coordinates a Ca^2+^ ion which might play a role in an earlier phase of PQQ binding (Figure 1f). A total of 12 amino acids from PqqU interact with PQQ and originate from extracellular loops 8 and 7, the N-terminal plug domain, and the β-barrel (Figure 1g,h). In addition, five water molecules directly coordinating PQQ were resolved, due to the high resolution of the cryo-EM data. Three arginine residues R304, R365, and R536 interact with the carboxylic groups of PQQ, significantly contributing to the binding of PQQ by charge attraction. A plug-domain-located glutamine Q84 interacts with the two carbonyl groups of the central aromatic ring of PQQ, which may be important during the release of PQQ. While the plug domain tyrosine Y99 forms a π–π stacking interaction with the aromatic ring of PQQ. Q84, Y99, and Q121 may tether PQQ to the plug domain, facilitating its entry into the periplasm when the transporter is energised by TonB, which leads to displacement of the plug domain and formation of the transmembrane transport pore ^22-26^.

**Figure 2:**
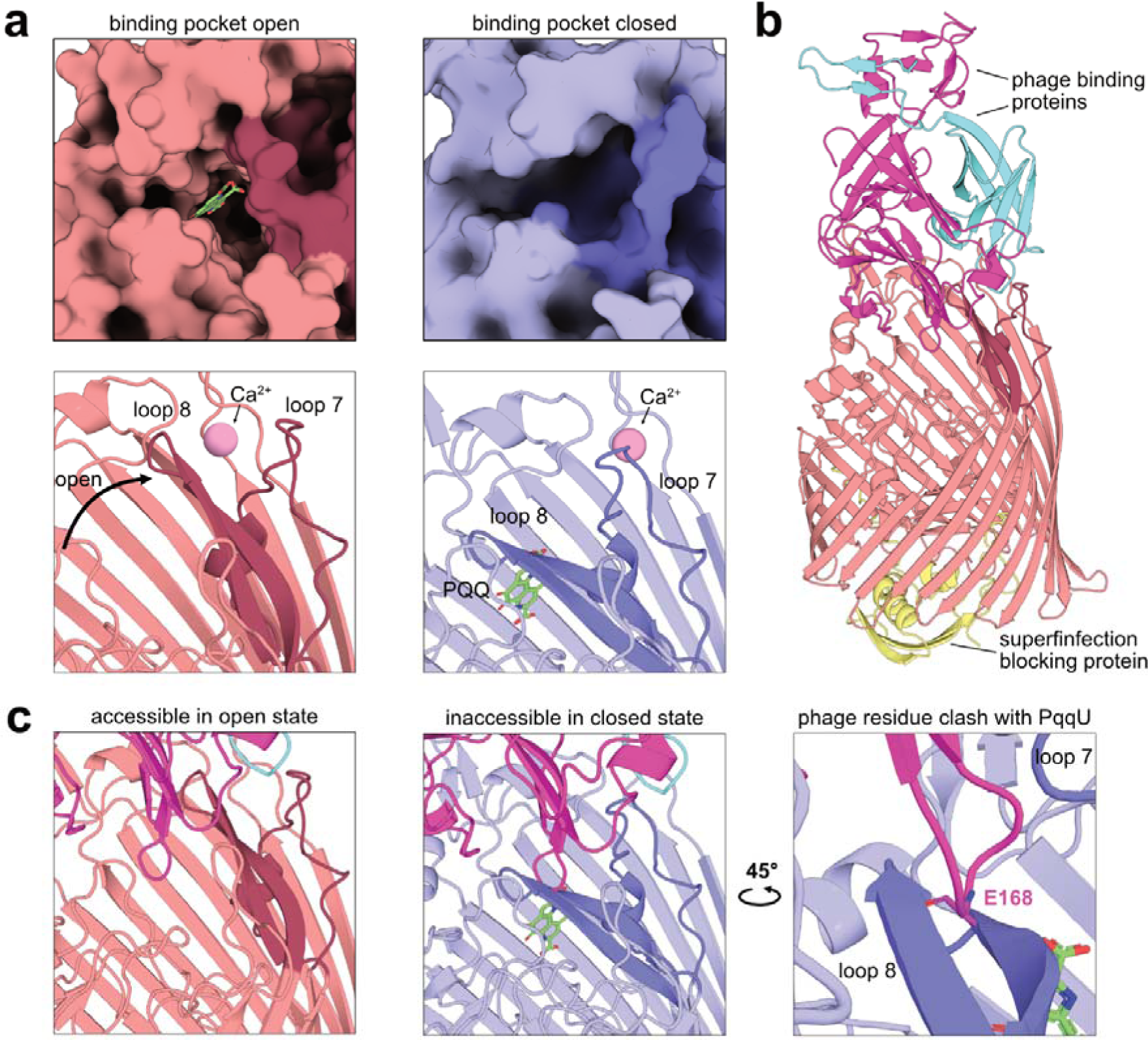
PQQ capture by PqqU prevents binding of the PqqU-targeting phage. **a** close-up comparison of the surfaces of PqqU in open (red) and closed (blue) states. PQQ was modelled into the apo PqqU structure, based on the position in the bound structure, to visualize that the binding pocket is open and accessible, whereas PQQ is hidden by loop 8 in the closed state. The close-up cartoon of PqqU in the closed conformational state (blue) compared to the open conformational state (red) visualizing the movement of loops 7 and 8 upon PQQ binding. **b**, AlphaFold2 prediction of PqqU with two IsaakIselin phage binding proteins (pink + cyan) and the putative superinfection blocking protein (yellow). **c**, the structure prediction indicates that the loops 7 and 8 must be in the conformationally open state to allow binding of the two phage proteins and otherwise clash with loop 8 in the closed state.

### Conformational changes in PqqU loops remodel the transporter to trap PQQ

In its unbound state, the substrate binding pocket of PqqU forms a deep cleft, which is open to the extracellular environment. In contrast, in the substrate-bound structure, PQQ is entirely enclosed within the substrate-binding pocket and is not accessible to the external environment (Figure 2a). This change in accessibility of the PqqU substrate binding pocket is largely mediated by major conformational changes in extracellular loops 7 and 8, which move inwards to form a barrier between the PQQ binding pocket and the external environment (Figure 2a, Movie S2). This change in the conformation of loops 7 and 8 is accompanied by a general compaction of PqqU around PQQ, as well as changes in the orientation of binding pocket sidechains to form the highly coordinated PqqU-PQQ complex (Figure 1g,h, Movie S3). Notably, Y99 and Q121 undergo a near 180° flip to interact with PQQ where Y99 forms the π–π interaction with the face to PQQ (Movie S3). These conformational changes are triggered by the binding of PQQ to the open state of PqqU, with our substrate-bound structure representing an intermediate state in the transport pathway, before substrate import facilitated by energy provided by interaction with TonB ^22-24,26^. The closure of the substrate binding pocket facilitates an airlock mechanism generally employed by TBDTs, where the PqqU transporter channel is not simultaneously open toward the periplasm and extracellular space, preventing the inadvertent import of deleterious compounds ^25^.

Previous studies discovered a family of bacteriophages that infect *E. coli* by utilising PqqU as their cell surface receptor and identified that PQQ can block infection by these phages ^16,27^. A 1000-fold lower PQQ concentration is required to block infection in an *E. coli ΔtonB* strain, in which PqqU cannot be activated for import ^16^. Given the high affinity of the PqqU-PQQ interaction and the closure of the binding pocket upon substrate binding, PqqU in the *ΔtonB* strain likely remains in the closed PQQ bound state indefinitely, suggesting that the closed state of PqqU identified by our structure is not competent for phage binding. To determine the structural basis for the observation, we used AlphaFold2 to predict the complex of PqqU and three proteins indicated as putative receptor binding proteins encoded by the PqqU targeting phage IsaakIselin ^27-29^. Strikingly, AlphaFold2 predicted a complex between PqqU and all three phage proteins with high confidence (Figure 2b, Figure S5, Supplemental Data S1). Two of the phage proteins (QXV80560, QXV80561) dimerise to form a complex with the extracellular face of PqqU, together constituting the phage receptor-binding domain. In contrast, the third protein (QXV80559) forms a complex with the intracellular side of PqqU and likely constitutes a superinfection prevention factor analogous to that of Llp from T5 phage ^30^. Interestingly, Foldseek analysis of the QXV80559 structural model reveals it is structurally homologous to the C-terminus of TonB, which binds TBDTs and provides the mechanical force required to drive substrate import ^22,31^. This indicates that the IsaakIselin phage has repurposed this domain of TonB to inactivate PqqU and block superinfection. In the AlphaFold2 model, the phage receptor binding domain binds to PqqU in the open state (Figure 2c), significant clashes occur between loops 7 and 8 of PqqU and the phage receptor if the closed PQQ-bound structure of PqqU is superimposed with this model. These clashes would likely preclude binding, explaining the observed resistance to the phage when PQQ is present (Figure 2c) ^16^. Taken together these data indicate that the closed PQQ-bound state we observe in our structure is likely irreversible without energy input from TonB, which supports our proposed airlock-style gating mechanism for PQQ import through PqqU.

### Electrostatic and π–π interactions are critical for PQQ transport by PqqU

To test the relative importance of residues in the PQQ binding site for substrate import, we generated PqqU variants where these residues were either mutated to alanine or in the case of K274, R304, R365, and R536 were charge swapped to glutamate (Figure 3). An inducible plasmid encoding these variants was transformed into a PTS-deficient *E. coli* strain also lacking PqqU (*E. coli ΔptsΔpqqU*). This strain requires the PQQ-dependent enzyme Gcd to grow when glucose is its sole energy and carbon source, which makes PqqU essential when external PQQ concentrations are low (<100 nM) ^16^. Under these conditions, after a significant lag phase (∼10 h) cell growth of both wildtype and mutant PqqU-expressing cells was measured by optical density. In the absence of exogenous PQQ, *E. coli ΔptsΔpqqU* complemented with WT PqqU did not exhibit significant growth, whereas, in the presence of 10 nM PQQ, the strain grew to a final OD of ∼0.8, with a doubling rate of approximately 4 hours. The exponential growth rate of strains complemented with several PqqU binding site variants (Y481A, Q308A, Q84A, Q310A, N447E) was comparable to wildtype, indicating that these residues are not critical for PQQ uptake by themselves. Conversely, a longer lag phase and significantly slower growth were observed for strains containing PqqU variants K274E, R304E, and R365E (Figure 3a,b). These residues are located in the β-barrel portion of the binding pocket and form salt-bridge interactions with PQQ, which indicates they are likely crucial in capturing the substrate. While PqqU variants R536E and Y99A did not exhibit a significantly slower growth rate during exponential phase, these variants exhibited a longer lag phase, indicating they are important for PQQ binding or transport (Figure 3a,b). Y99A is located in the plug domain and forms a direct π–π stacking with the polycyclic PQQ, stabilising the cofactor during binding and possible import. These data indicate that while this interaction is important for PqqU function it is not as critical for function as the complementary charge-charge interactions with the PQQ carboxylic acid groups. In summary, this section identified key residues in the PqqU binding site for PQQ import, providing mechanistic insight into transporter function, and providing a blueprint for identifying PQQ transporters in other bacteria.

**Figure 3:**
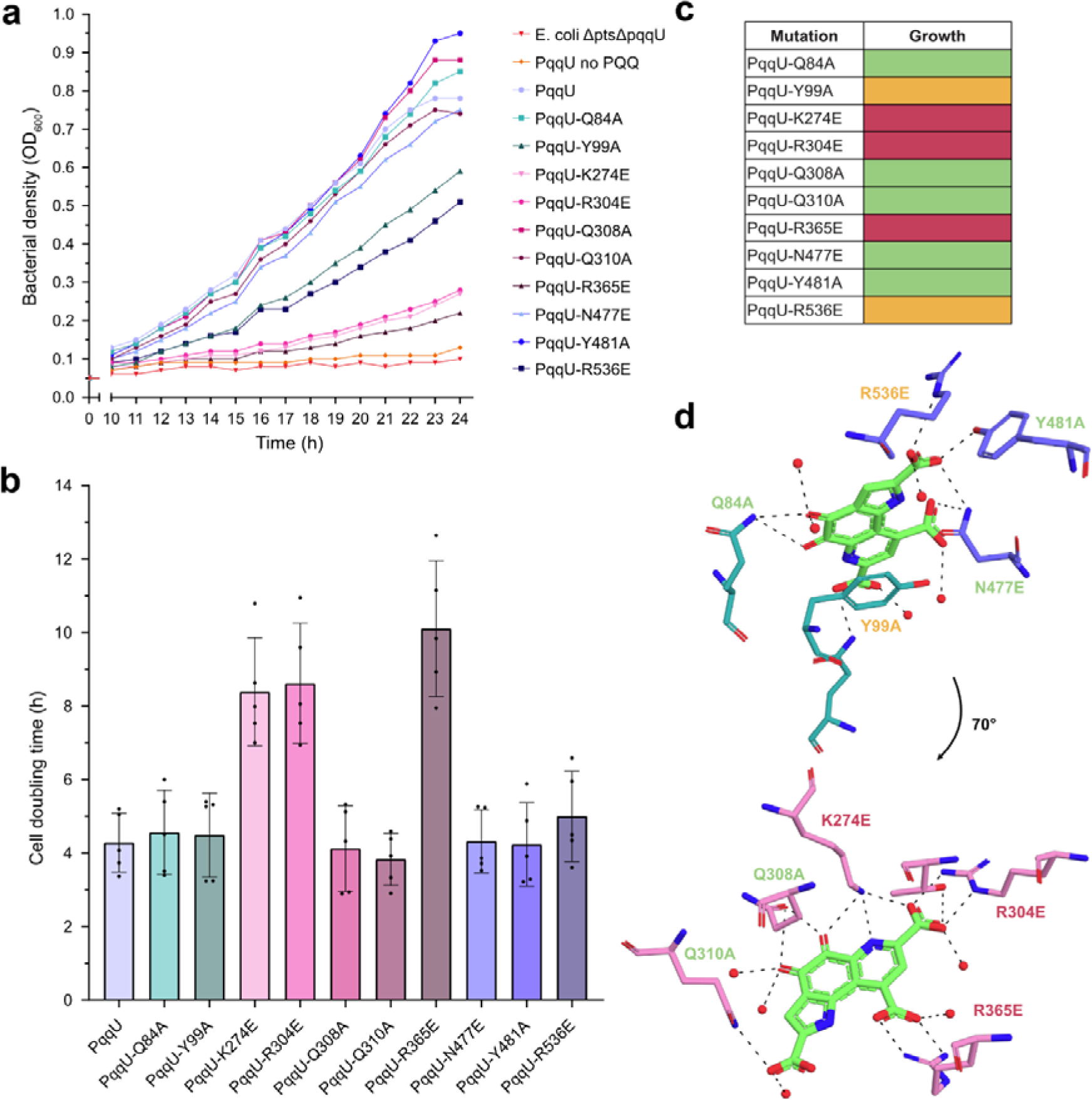
Active site mutations of PqqU reduce the ability to perform EDP glycolysis. **a**, a representative growth curve of *E. coli* ΔptsΔpqqU cultures grown in M63 minimal media containing 0.1 % glucose and supplemented with 10 nM PQQ unless indicated. The cells were transformed with plasmids containing mutated active site variants of pqqU. **b**, the doubling time of cells was individually calculated from the exponential growth phase of each culture (n = 5, biological replicates). Statistical differences between PqqU and mutated PqqU were determined by performing paired t-tests (ns = not significant, * = p value <0.05, ** = p-value <0.01). **c**, an overview of the effect of active site mutations on PQQ-dependent growth: No effect (green), delayed growth (orange), and growth defect (red). **d**, rotational view of PQQ with the 11 active site residues. The mutations and their effect on culture growth are coloured based on severity.

### PqqU is present in diverse Gram-negative bacteria

TonB-dependent transporters import diverse substrates and are highly variable in amino acid sequence and outer loop structure, even between transporters targeting the same substrate ^22,32-34^. As a result, it is difficult to definitively determine substrate specificity based on phylogenetic relationships and global sequence identity alone. To overcome this and identify the presence of PqqU in Gram-negative bacteria, we utilised the key conserved PQQ binding residues Y99, R304 and R365 as a molecular signature for PqqU, to identify the presence of genes encoding PqqU in publicly available genomes and metagenome-assembled genomes (MAGs) (Figure 3d). As Y99 is responsible for π–π interactions with PQQ, we also accepted phenylalanine and tryptophan in this position, as they will also satisfy this interaction. After genome dereplication, PqqU was identified in 1,861 diverse Gram-negative bacterial isolate genomes and MAGs, representing 22 phyla. We subsequently analysed the genomes of these PqqU-producing isolates for the genes responsible for PQQ biosynthesis (PqqA-PqqF) and PQQ-dependent dehydrogenases, to gain insight into the role of PqqU in these organisms (Figure 4a, Figure S6, Supplemental Data S2 and S3).

**Figure 4:**
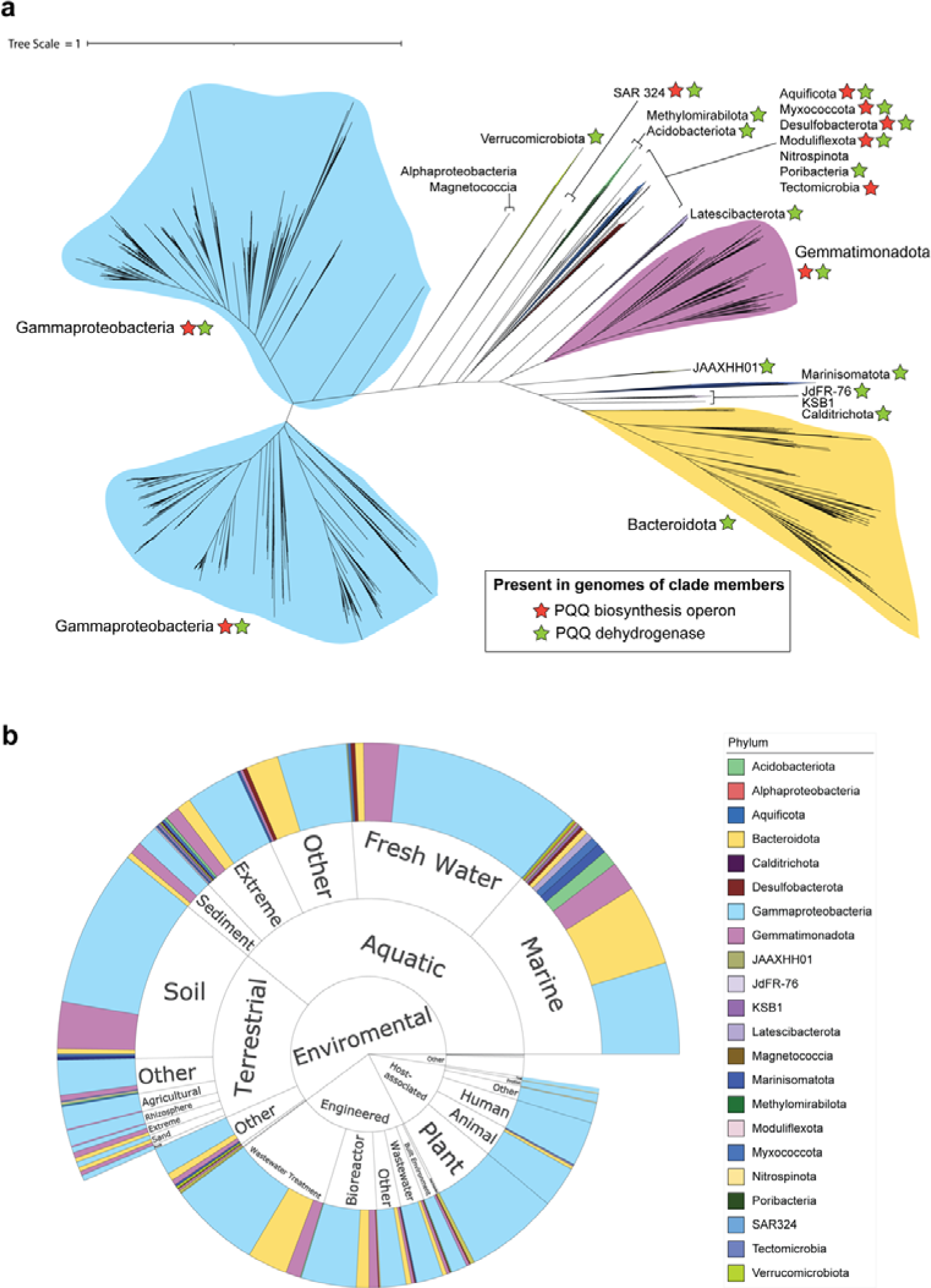
The phylogeny and origin of PqqU-containing bacteria. **a,** a phylogenetic tree of bacteria encoding PqqU identified by our homology search of bacterial genomes. The tree was constructed based on 16 ribosomal proteins. The classes of bacteria encoding the various clades of PqqU are shown, as well as the presence of genes encoding PQQ biosynthetic capacity or quinonoproteins (PQQ-dependent dehydrogenases) in at least some of the members of that clade. See Figure S4 for a more detailed view of this tree. ***b***, A sunburst plot showing the environment of origin/isolation for bacteria encoding PqqU.

**Figure 5:**
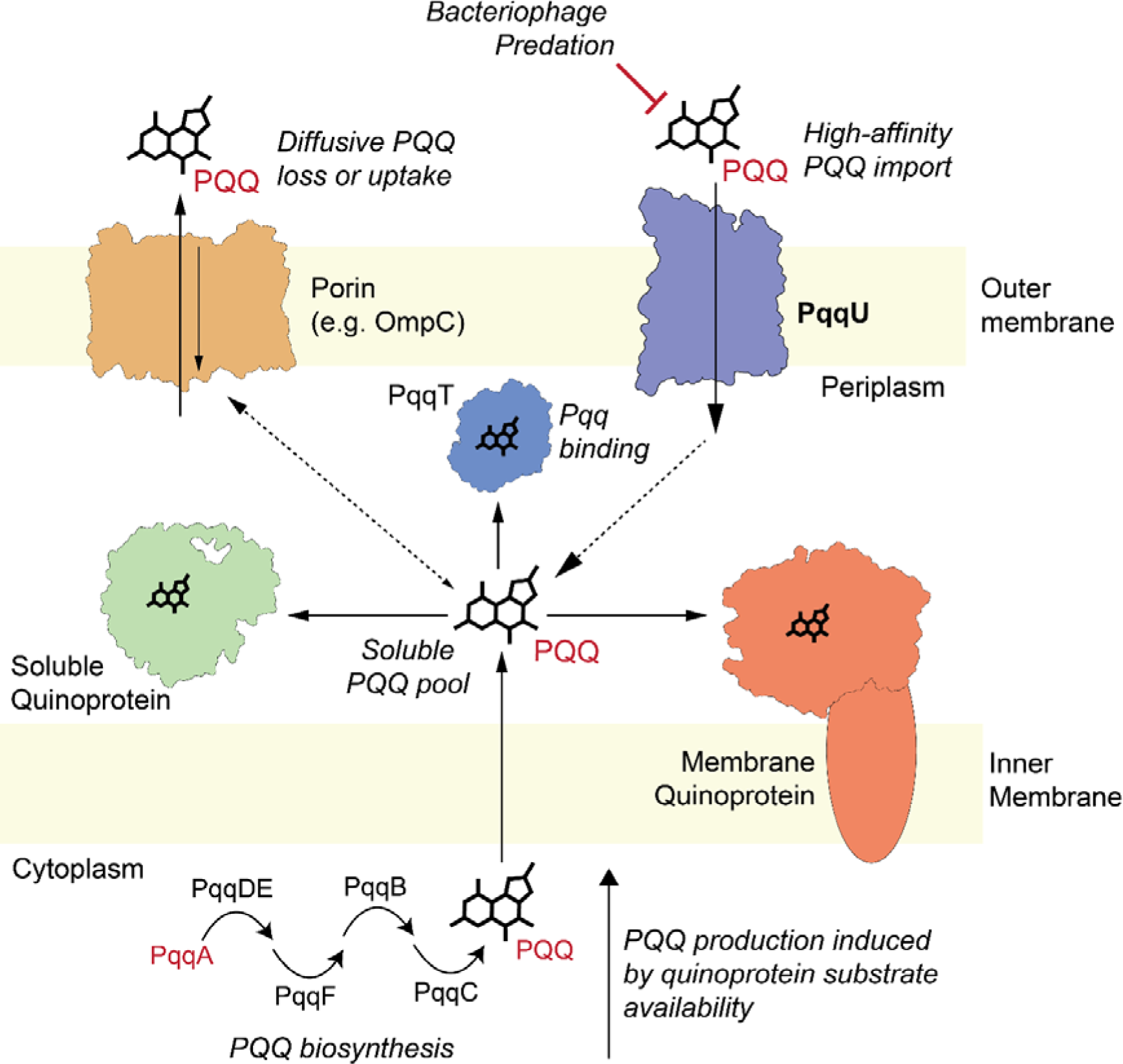
A model of the different strategies employed by Gram-negative bacteria for obtaining PQQ for use in quinoproteins. This diagram is a composite of the different strategies employed by Gram-negative bacteria to obtain PQQ. Based on our genomic analysis some strains employ both PQQ biosynthesis and scavenging by PqqU to obtain PQQ while others rely on one of these strategies.

Most bacteria encoding PqqU were Gammaproteobacteria (n = 1346), which is present in 21 orders of this class. This is consistent with the abundance of genomes belonging to this class in available databases. Other notable PqqU-producing classes include Bacteroidota (n = 206), Gemmatimonadetes (n = 183), Acidobacteria (n=17), Latescibacterota (n=14), and Marinisomatota (n=11) (Figure 4a, Figure S6). Interestingly, PqqU was largely absent from Alphaproteobacteria (n=1), even though these bacteria commonly contain PQQ-dependent quinoproteins and produce PQQ ^7,35,36^, indicating that they are likely primarily PQQ producers rather than scavengers. However, this does not appear to be the predominant strategy adopted by PqqU producing bacteria, as out of the 22 phyla with members encoding PqqU only 8 contain the machinery for PQQ biosynthesis. In total, the genes for PQQ biosynthesis could be identified in only 23 % (n = 465) of PqqU producers and were absent from Bacteroidota (n=206), Acidobacteria (n=17), Latescibacterota (n=14), Marinisomatota (n=11) and Verrucomicrobia (n=5), suggesting bacteria from these phyla rely exclusively on PQQ scavenging. When both PqqU and the PQQ biosynthesis operon were present, they generally did not show synteny, suggesting they are regulated independently. However, we observed four Burkholderiales genomes showing synteny and the same gene orientation for these genes indicating co-regulation occurs in some organisms (Figure S7). Overall this analysis indicates that the scavenging of PQQ from the environment at low concentrations using PqqU is a widespread strategy for the utilisation of this cofactor.

PQQ is essential for the function of a diverse family of quinoprotein dehydrogenases, including lanthanide-dependent methanol dehydrogenases (XoxF clades), which are globally abundant and highly expressed in both soil and marine environments ^10,37^. While most PqqU producers we identified encode at least one predicted PQQ-dependent dehydrogenase (67 %), no PQQ-dependent enzymes were identified in members of some Gammaproteobacterial orders/genera (Nitrosomonadaceae (n = 40), Ectothiorhodospirales (n = 8), Arenimonas (n = 7)) and bacterial phyla Nitrospinota (n = 5) and Ignavibacteriota (n = 4). These bacteria also do not synthesise PQQ, suggesting an alternative role for the cofactor in these bacteria, possibly via a PQQ-dependent enzyme not identified in our search, or as an antioxidant.

PqqU-producing bacteria are ecologically diverse, occurring in a wide range of aquatic, terrestrial, host-associated, built environments, and extreme conditions (high and low pH and temperature). This indicates that PQQ is ubiquitous, chemically, and thermally stable and an important nutrient that contributes to the adaptability of bacteria to various environments (Figure 4b). Gammaproteobacterial PqqU producers were found in all environment categories and are frequently capable of both producing and scavenging PQQ, suggesting they play a key role in determining environmental PQQ concentrations. Acidobacteria and Bacteroidota phyla are ubiquitous and abundant members of soil microbiomes and are known to utilise quinoproteins^38,39^. These phyla are well represented in public databases but despite this, we only identified PqqU producers in class Vicinamibacteria (Acidobacteria) and classes Ignavibacteria, Rhodothermia and Bacteroidia (Bacteroidota) that were primarily isolated from aquatic environments (Figure 4b). That members of these phyla that do not encode PqqU may rely on endogenous PQQ production or that soil PQQ concentrations are high enough to satisfy their requirements by diffusion. Plant growth-promoting bacteria are often found to produce PQQ. Under phosphate limiting conditions they induce PQQ production which is needed for a PQQ-dependent glucose dehydrogenase ^40^. The secreted gluconate solubilizes phosphate minerals and furthers bacterial and plant growth ^41^.

PqqU-producing bacteria also include significant human, animal, and plant pathogens including multidrug-resistant *Pseudomonas aeruginosa*, *Klebsiella pneumoniae, Salmonella* sp. and *E. coli* (Figure 4b and Figure S6). In members lacking PQQ biosynthesis machinery, PqqU may contribute to bacterial virulence by enabling the metabolic versatility provided by quinoproteins^1^, which would be beneficial in nutrient-limited environments inside the host, or in environmental reservoirs. In *Salmonella* Serovar Typhi, PqqU (YncD) was reported to be important for virulence in a mouse intraperitoneal infection model. The attenuated *ΔpqqU* strain was used for vaccination and protected against subsequent infection with the wildtype strain ^42^. Subsequent work indicates that vaccination with recombinant PqqU from *S.* Typhi also protects against its infection in a mouse model. However, this study also suggested that PqqU is involved in iron uptake, which this current study and previous work strongly indicate is not the case ^43^. If PQQ scavenging by PqqU is important for virulence, it suggests that it is an important nutrient for bacteria even within animals or the human host. However, considering animals do not synthesize PQQ its presence in the body must result from dietary intake or synthesis by the gut microbiome ^44^. This would likely make PQQ more abundant in the gastrointestinal tract, although the high affinity of PqqU could make it compatible with scavenging trace PQQ present in human tissues. Further work is required to shed light on the possible role of bacterial PQQ import during infection. Finally, the presence of PqqU in many pathogens suggests it has potential as a cell surface receptor for engineered bacteriophage therapies, for example, the PqqU-dependent phage Isaaklselin in *E. coli* ^16,45^.

## CONCLUSIONS

This work demonstrates that PqqU from Gram-negative bacteria binds PQQ with high affinity, facilitating the import of low concentrations of this cofactor from the environment. By resolving the CryoEM structure of the 75 kDa PqqU in complex with PQQ at the high resolution of 1.99 Å, we show that PQQ is highly coordinated within an internal cavity of PqqU, which is formed by significant conformational changes of the transporter’s extracellular loops upon PQQ binding. This conformational change traps PQQ inside the transporter, blocks binding of PqqU-dependent bacteriophage, and likely facilitates an airlock-style import mechanism that prevents the uptake of unwanted compounds. This work also confirms that PqqU transports a non-metal-containing substrate, which is unusual for a TBDT and underpins the importance of PQQ as an external nutrient for Gram-negative bacteria ^22^. Utilising the knowledge of the PQQ binding site revealed by our CryoEM structure and functional analysis, we show that PqqU facilitates PQQ import in Gram-negative bacteria from 22 phyla, which are present in diverse environments. This indicates that PQQ is an important externally derived nutrient and a ubiquitous common good of diverse microbiomes. The presence of PqqU and quinoproteins in many bacteria that lack the machinery for PQQ biosynthesis indicates that PQQ scavenging is a widespread strategy for satisfying metabolic requirements for this cofactor, which may be used in conjunction with PQQ production, and sequestration to prevent its loss from the cell via diffusion (Figure 6).

## MATERIALS AND METHODS

### Bacterial strains and growth conditions

For cloning, *E. coli* DH5α were maintained on LB agar plates and broth (10 g/l Tryptone, 5 g/l Yeast extract, 5 g/l NaCl, 15 g/l agar) with 100 µg/ml Ampicillin where appropriate. Cultures were incubated at 37°C with orbital shaking at 180 rpm.

For expression of 10xHis-PqqU, *E. coli* BL21 (DE3) C41 were transformed with the plasmid pET-20b-pqqU. A starter culture was inoculated with 90% of the transformation and grown in LB broth (100 µg/ml Ampicillin) overnight at 37°C with rotary shaking at 180 rpm. The remaining 10% were plated as contamination control. Main cultures were inoculated with the starter culture to start OD_600_ 0.1 and grown until OD_600_ 1 in TB broth (12 g/l Tryptone, 24 g/l Yeast extract, 12.3 g/l K_2_HPO_4_, 2.52 g/l KH_2_PO_4_) at 37°C at 180 rpm. Cultures were cooled to 4°C after which the expression was induced with 0.3 mM IPTG. Cultures were grown for pqqU expression for ∼18 h at 24°C at 180 rpm.

For complementation experiments, *E. coli* BW25113 ΔptsΔpqqU starter cultures were grown in LB broth containing 0.05% L-arabinose and 100 µg/ml Ampicillin where appropriate. Complementation main cultures were grown in M63 media (14.13 mM (NH_4_)_2_SO_4_, 38.95 mM KH_2_PO_4_, 1 mM MgSO_4_, 3.3 µM FeSO_4_, 0.05% arabinose, pH 7) and where appropriate 0.1% glucose and 10 nM pyrroloquinoline quinone. Main cultures were inoculated with start OD_600_ 0.05 after resuspending starter culture cell pellet in M63 media + 0.05% L-arabinose.

### Generation of PqqU mutants by site-directed mutagenesis

Active site mutations of PqqU were generated from the plasmid template pBAD24-pqqU using the Q5® Site-Directed Mutagenesis Kit (New England BioLabs) according to the manufacturer’s instructions. Mutations were confirmed by nanopore sequencing (Primordium Labs). See Table S3 for primer sequences.

### Complementation growth experiments of PqqU mutants

Growth curves were generated from a total of 4 biological and 2 technical replicates (5 growth curves in total). Starter cultures were prepared as previously above, grown overnight, and kept on ice during the day. 2 h before inoculation of the main cultures, starter cultures were centrifuged at 4,500 xg for 10 min and resuspended in M63 media containing 0.05% L-arabinose. Complementation cultures were prepared as described above. Complementation cultures were grown in M63 media providing 0.1% glucose as an energy/carbon source and 10 nM PQQ as the cofactor for Gcd activity. Cells required a 10 h equilibration time before we could measure the growth for 14 h. Growth curves were plotted and analyzed with the GraphPad Prism 10.0.3 software.

### PqqU purification

PqqU (Formerly YncD) was purified from *E. coli* C41 containing pET-20b-pqqU on 8 occasions from either 8 l or 16 l of LB media per purification. The yield of PqqU was 150 µg/l on average, depending on pellet size and which detergent was used for membrane solubilization. Cell pellets were resuspended in a final volume of 100 ml lysis buffer (50 mM Tris-HCl pH 7.4, 150 mM NaCl, 10 mM Imidazole, 2 mM MgCl_2_, 0.1 mg/ml Lysozyme, 0.05 mg/ml DNase, 1x protease inhibitor) and homogenized with a glass homogenizer before being subjected to cell lysis in a high-pressure cell disruptor. The lysate was centrifuged at 27,000 xg for 20 min at 4°C. The resulting supernatant was used for ultracentrifugation at 100,000 xg for 40 min at 4°C. The membrane pellet was resuspended in 40 ml of solubilization buffer (50 mM Tris-HCl pH 7.4, 150 mM NaCl, 10 mM Imidazole, 5% Elugent detergent) and homogenized in a glass homogenizer. Solubilization was done at room temperature with slow rotary shaking. The sample was centrifuged at 27,000 xg for 20 min and the resulting supernatant was loaded 2x on a HisTrap HP 5ml affinity column. After extensive washing with Wash buffer (50 mM Tris-HCl pH 7.4, 500 mM NaCl, 10 mM Imidazole, 0.03% DDM or 0.025% LMNG), PqqU was eluted with a step gradient in elution buffer (50 mM Tris-HCl pH 7.4, 150 mM NaCl, 1 M Imidazole, 0.03% DDM or 0.025% LMNG). When purified in n-dodecyl-β-D-maltoside (DDM), PqqU eluted in 250 mM Imidazole, while it eluted in 500 mM Imidazole when purified in Lauryl Maltose Neopentyl Glycol (LMNG). Elution fractions were analyzed via SDS-PAGE and pooled and concentrated to a volume of 5 ml in an Amicron Ultra-15 Centrifugal Filter 30 kDA MWCO Millipore. Size exclusion chromatography was done with a Superdex HiLoad 200pg 26/600 in size exclusion buffer (50 mM Tris-HCl pH 7.4, 150 mM NaCl, 0.03% DDM or 0.025% LMNG). PqqU eluted in a single peak at a retention volume of 140-180 ml when purified in DDM, and in a double peak at a retention volume of 140-180 ml. PqqU could be concentrated to up to 15 mg/ml without significant loss of protein in an Amicron Ultra-15 Centrifugal Filter 100 kDA MWCO Millipore and stored at -80°C without glycerol. Before cryo-EM and ITC, 20-100 µl of PqqU (Purified in LMNG) were diluted in 50 ml size exclusion buffer without detergent and re-concentrated to remove excess detergent.

### SDS-PAGE analysis

Protein samples were run on a Bolt 4-12% SDS-PAGE gel according to the manufacturer’s instructions. Gel staining was done with AcquaStain Protein Gel Stein (Bulldog) and subsequently washed in distilled H_2_O. Protein samples were generally loaded by constant volume.

### Cryo-EM imaging and dataprocessing

Cryo-EM imaging was performed on PqqU which was initially purified in DDM and then detergent exchanged to LMNG with subsequent removal of excess LMNG as described above.

#### Grid preparation

UltrAuFoil gold grids (Quantifoil GmbH)^46^ were glow discharged at 30 mA for 60 s (PELCO easiGlow^TM^) in atmosphere. A 3 µl of the sample was applied at a concentration of 43 µM PqqU (3.3 mg/ml) with 910 µM PQQ (Dissolved in H_2_O) to the glow discharged grids. The excess protein solution was blotted off using a blot force of 0 and a blot time of 3 seconds in a Mark IV vitrobot (Thermo Fischer Scientific) which was maintained at 100% humidity and 4°C.

#### Imaging

Imaging was performed on a G1-Titan Krios (Thermofisher) with Schotky FEG operated at 300 kV and equipped with Gatan K3 mounted post a Gatan BioQuantumn energy filter. The fast data collection strategy in EPU (Thermo Fischer Scientific) enabled with aberration-free image-shift (AFIS) alignments was used for automated data collection. Zero-loss energy-filtered images were acquired at a nominal magnification of 105,000X using a narrow energy filter slit width of 10eV. K3 was operated in counting CDS mode with a pixel size of 0.82 Å. Movies were collected with a total dose of 60 e^−^ Å^−2^ accumulated over 5.22 s exposure time at a dose rate of 8.518 e^−^ pixel^−1^ s^−1^ fractionated into 60 frames. A total of 8744 movies were recorded.

#### Image processing

The resultant movies were dose-weighted and motion-corrected using UCSF Motioncor2 (version: 1.6.3)^47^ to output both dose-weighted and non-dose-weighted averages. The non-dose-weighted averages were used for estimating contrast transfer function (CTF) parameters using CTFFIND 4.1.8^48,4948,4949,5049^, using RELION 4.0.1^50^ wrapper. Particle picking was performed using Gautomatch(0.53) (https://www.mrc-lmb.cam.ac.uk/kzhang/) on the full dataset using a blob diameter of 80 Å. This resulted in 2,112,379 particles from resultant coordinates which were extracted and binned 4X to a box size of 64 pixels. These particles were imported to cryoSPARC^51^ (version:4.2.0) and subjected to 2D classification to retain 482,616 particles. Ab initio classification with class similarity of 0 and 3 classes was performed to result in 258,422 particles giving a consensus ab-initio model that had features of micelle-associated PqqU. 3D refinement in cryoSPARC 4.2.0 was performed to further centre the particles. The coordinates were then exported back to RELION using pyem v0.5^52^ and re-extracted unbinned in RELION 4.0.1 centred on the refined coordinates. The particles were reimported back to cryoSPARC 4.2.0. One round of 2D classification resulted in retaining 252,169 particles for further processing. A round of homogenous refinement followed by 2 rounds of non-uniform(NU) refinement^53^ with defocus, tilt and trefoil refinement^53,54^ resulted in a 2.69 Å map. These particles were then subjected to Bayesian Polishing^55^ in RELION 4.0.1 and the resultant polished particles were then subjected to a round of homogenous refinement followed by 2 rounds of NU refinement to yield 2.49 Å map. Further, baited heterogenous refinement with 2 classes with one of them being a noisy reconstruction was used to filter away junk particles. The filtered particles were then subjected to further NU refinement and heterogenous refinement to retain 148,172 particles resulting in 2.29 Å resolution map. To push the resolution further, these particles were used to train TOPAZ particle picker^56^ on the full dataset. The trained model was then able to pick a total of 6,488,281 particles. These were then subjected to the same processing workflow as above. A round of 2D classification resulted in retaining 1,179,837 particles. These were then subjected to a round of homogenous refinement followed by baited heterogenous refinement to yield 872,884 particles. Further NU refinement on the resultant particles along with CTF refinement yielded a 2.18 Å resolution map. These particles were then subjected to Bayesian Polishing in RELION 4.0.1 and were further subjected to NU refinement with CTF refinement to yield 2.06 Å resolution map. A final round of local refinement with a micelle-subtracted map (generated using UCSF Chimera (version 1.16)^38^) derived mask resulted in the final 1.99 Å resolution map (gold standard FSC 0.143 criteria).

### PqqU + PQQ model building and visualization

The apo crystal structure of PqqU was used for docking into the 1.99 Å resolution map of PqqU using ChimeraX 1.6.1. Model building was done using WinCoot CCP4 0.9.8.1. The structural coordinates for PQQ were extracted from PDB entry 7WMK and used to fit in the experimental maps. PHENIX was used for real-space refinement and DOUSE was used for initial identification of H_2_O but had to be heavily corrected. Model quality was assessed and confirmed using the PDB validation web tool. Images were created using ChimeraX and PyMOL.

### Isothermal titration calorimetry

PqqU for isothermal titration calorimetry was purified in LMNG as described previously. On the day before the ITC experiment, the detergent-removed sample was concentrated to ∼150 µl before dialysis in a 3.5 kDa cut-off dialysis bag (Spectra/Por Dialysis Membrane 3.5 kD) in 2 l of dialysis buffer (50 mM Tris-HCl pH 7.4, 150 mM NaCl) at 4°C overnight on a magnetic stirrer on low speed. Dialyzed PqqU was recovered and centrifuged at 13,000 xg for 10 min at 4°C. The PqqU concentration was measured after centrifugation and PQQ (Methoxatin disodium salt) was dissolved in the same overnight dialysis buffer to a stock concentration of 5 mM PQQ. ITC was performed using the Malvern MicroCal PEAQ ITC. The 3 technical replicates and controls were performed from the same purified PqqU sample. 100 µM PQQ was titrated into 20 µM PqqU over 40 minutes in 19 injections of 2 µl. Graphs and figures were generated from base-line corrected data in the GraphPad Prism 10.0.3 software.

### Bioinformatic Analysis

The NCBI NR GenBank release 254 (February 15 2023) database was queried using blastp v2.14.1+ for PqqU (NP_415968.1) via the command line with an e-value cutoff of 1E-5. Sequences were also retrieved from the UniProt reference proteome database using an e-value cutoff of 1E-5. Retrieved sequences were aligned using FAMSA (Fast and Accurate Multiple Sequence Alignment) ver. 1.2.5 ^57^ with default parameters and trimmed using trimal v1.4.1 and the -gappyout option ^58^. Sequences were manually verified based on the presence of conserved PqqU substrate binding residues Tyr^99^ (or Phe^99^ or Trp^99^), Arg^304^ and Arg^365^ using Geneious ^59^. BioSample ID’s from verified PqqU homologs were used to retrieve genomes from the BV-BGC database ^54^. To account for any database inconsistencies, genomes were again searched and manually verified to contain PqqU homologs before being dereplicated using dRep 3.4.2 with default settings ^60^.

Genomes were searched for the PQQ biosynthesis operon (PqqA, PqqB, PqqC, PqqD, PqqE, PqqF) from *Pseudomonas aeruginosa* using cblaster v1.3.18 ^61^. PqqU (NP_415968.1) was also used as a query to determine if it is colocalized with the PQQ biosynthesis operon. The colocalization and orientation of the genes were visualised using clinker v0.0. ^62^.

Pyrroloquinoline quinone-dependent dehydrogenases were identified using a HMM profile taken from Diamond et al. 2019 ^39^. Pyrroloquinoline quinone protein domains were predicted in the retrieved sequences by using hmm search (version 3.3) ^63^ to search against the protein family database (version 36.0) ^64^ for PQQ pfams PF01011, PF13360, and PF13570 with an e-value cutoff of 1E-5.

A ribosomal protein tree was built using a concatenation of 16 ribosomal proteins identified in each genome using GOOSOS.py (https://github.com/jwestrob/GOOSOS). The following HMMs were used: Ribosomal_L2 (K02886), Ribosomal_L3 (K02906), Ribosomal_L4 (K02926), Ribosomal_L5 (K02931), Ribosomal_L6 (K02933), Ribosomal_L14 (K02874), Ribosomal_L15 (K02876), Ribosomal_L16 (K02878), Ribosomal_L18 (K02881), Ribosomal_L22 (K02890), Ribosomal_L24 (K02895), Ribosomal_S3 (K02982), Ribosomal_S8 (K02994), Ribosomal_S10 (K02946), Ribosomal_S17 (K02961), and Ribosomal_S19 (K02965). Ribosomal S10 model PF00338 was also used for the identification of Chloroflexi. All trees were constructed and decorated using iTOL ^65^.

## Acknowledgements

We acknowledge the use of instruments and assistance at the Monash Ramaciotti Centre for Cryo-Electron Microscopy, a Node of Microscopy Australia. This work was supported by a NHMRC Emerging Leader Grant (APP1197376) (to R.G.), an ARC Discovery Project (DP230102150) (to R.G.), and ARC LIEF grants (LE200100045, LE120100090) for the Titan Krios Gatan K3 Camera and for the Titan Krios.

## Data Availability Statement

Cryo-EM maps and atomic models generated in this paper have been deposited in the Protein Data Bank (accession code 9C4O) and the Electron Microscopy Data Bank (accession codes EMD-45192). AlphaFold2 models and source data are provided with this paper.

## Author Contributions

R.G., F.M., and M.V. conceived and designed the experiments. F.M., H.V., and R.G. performed the experiments. F.M., R.G., H.V. and M.V. analysed the data. R.G. and K.H. contributed reagents/materials/analysis tools. R.G., F.M., H.V. and M.V. wrote the paper. All authors edited and approved the manuscript.

## SUPPLEMENTAL FIGURES

**Figure S1:**
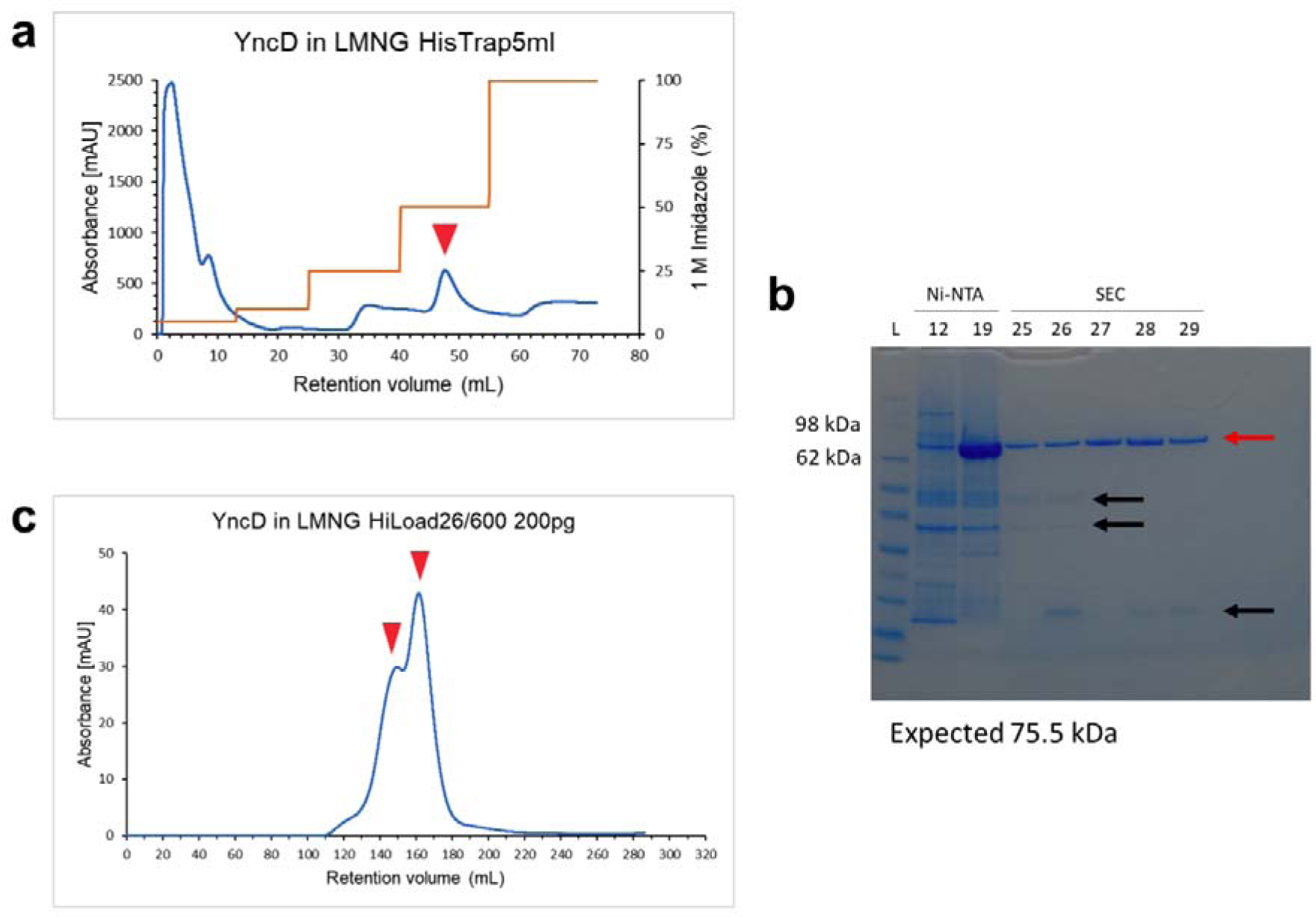
Purification of PqqU in LMNG. **A**, Elution profile (Blue) of PqqU from HisTrap HP 5-ml with a step gradient of Imidazole (Orange). The red arrow indicates the elution peak corresponding to PqqU. **B**, Elution profile of PqqU during size exclusion chromatography off a Superdex HiLoad 200pg 26/600 column. Red arrows indicate the double peak corresponding to PqqU when purified in the presence of LMNG. **C**, SDS-PAGE gel of affinity chromatography and size exclusion chromatography fractions corresponding to the observed double peak of the chromatogram. The red arrow indicates the band corresponding to PqqU (76 kDa) and the black arrows highlight minor contaminating proteins.

**Figure S2:**
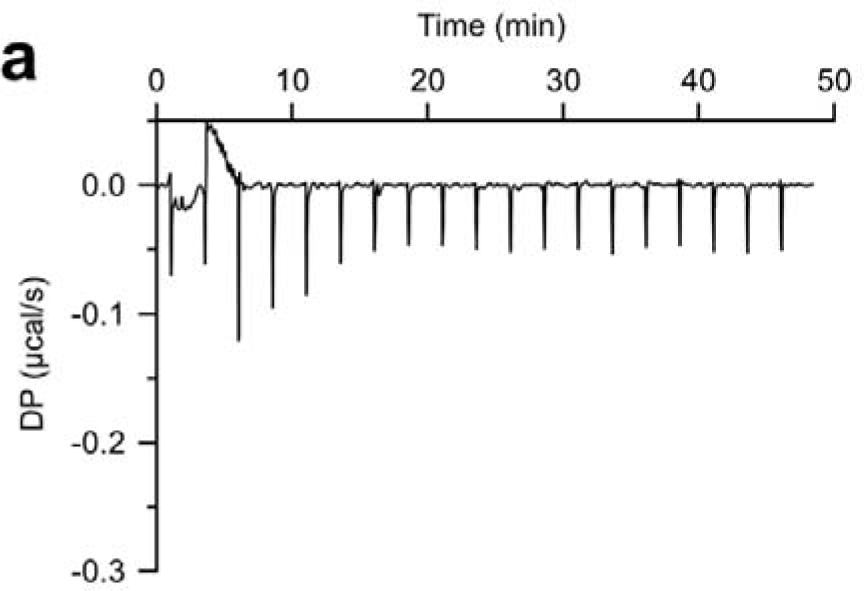
ITC buffer control injected into PqqU. **a**, buffer control of PqqU+PQQ isothermal titration calorimetry experiment. Dialysis buffer was injected into 20 µM PqqU over ∼50 minutes in 19 injections of 2 µl, the first injection is 0.4 µl. Experiments were conducted in triplicate on a Malvern MicroCal PEAQ-ITC and analysed with the accompanying software. Baseline corrected data were plotted with GraphPad Prism

**Figure S3:**
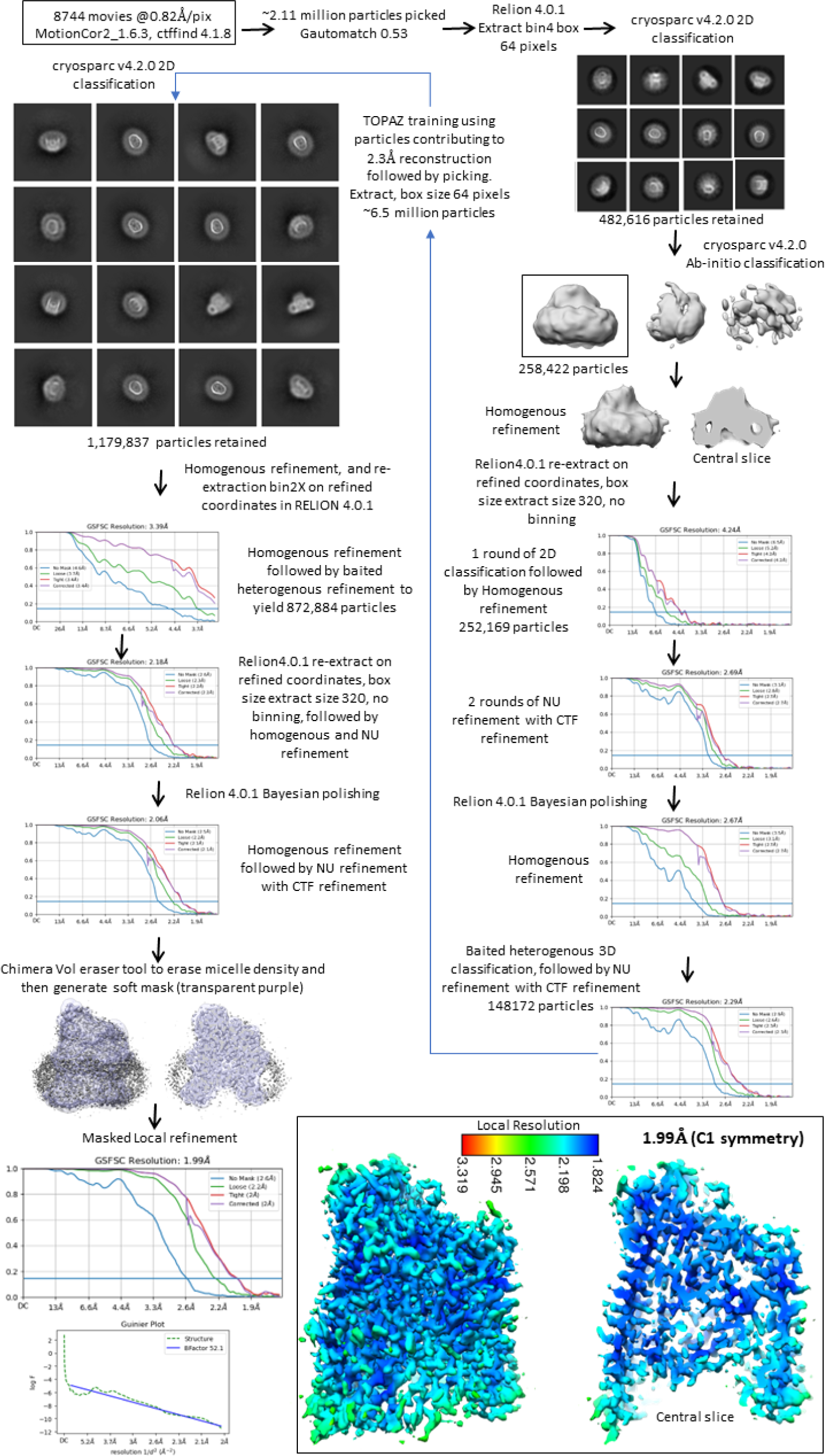
CryoEM data processing workflow for PqqU.

**Figure S4:**
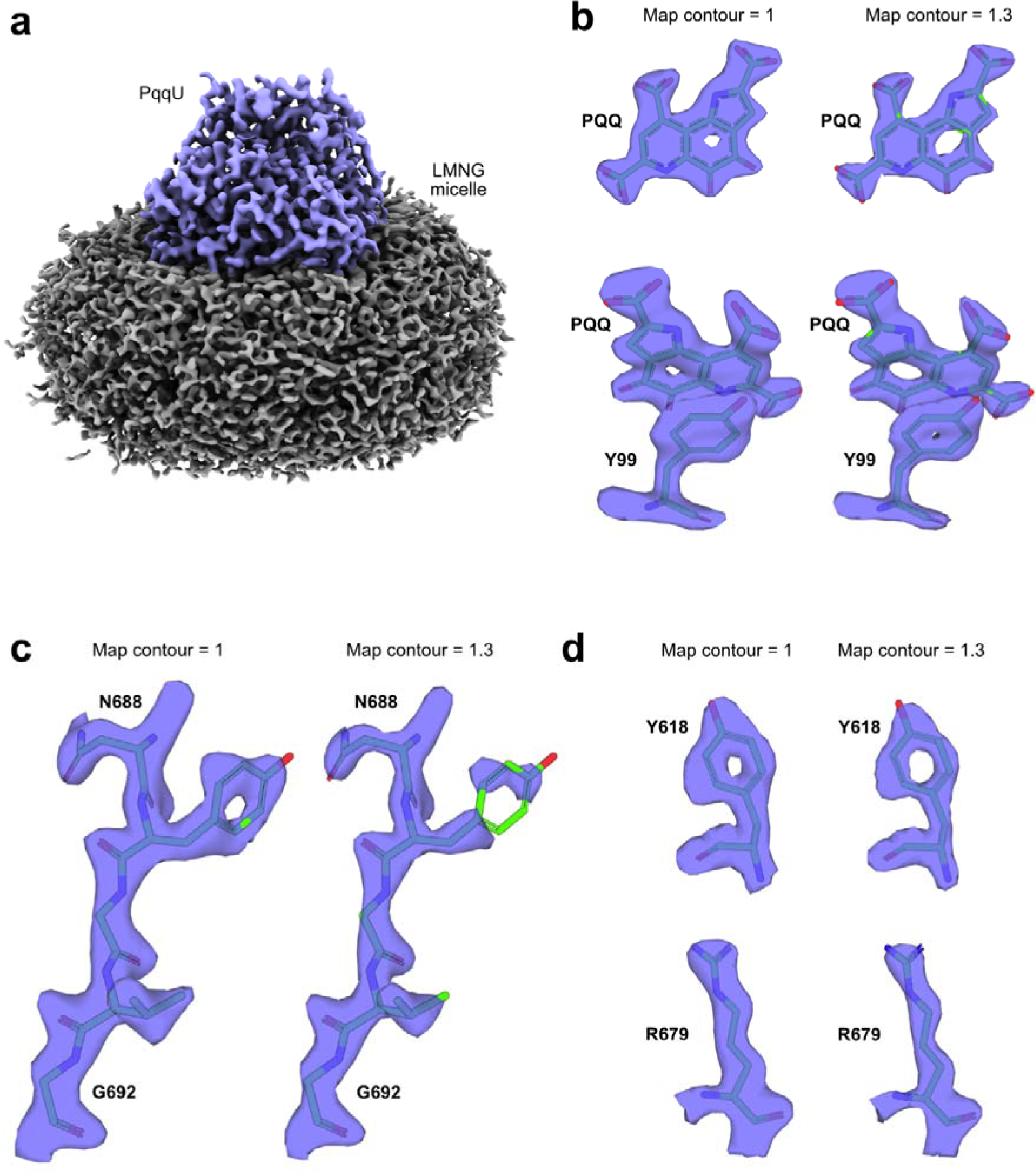
Additional structural images/analysis of PqqU and the PqqU-PQQ complex. **a**, Coulomb potential map of PqqU (Light-blue, contour 0.65) in its LMNG detergent micelle (Grey, contour 0.17). Map visualized with ChimeraX 1.7.1. **b-d**, close-up comparison of Coulomb potential maps of PQQ and selected residues at contours 1 and 1.3. Maps visualized with PyMOL.

**Figure S5:**
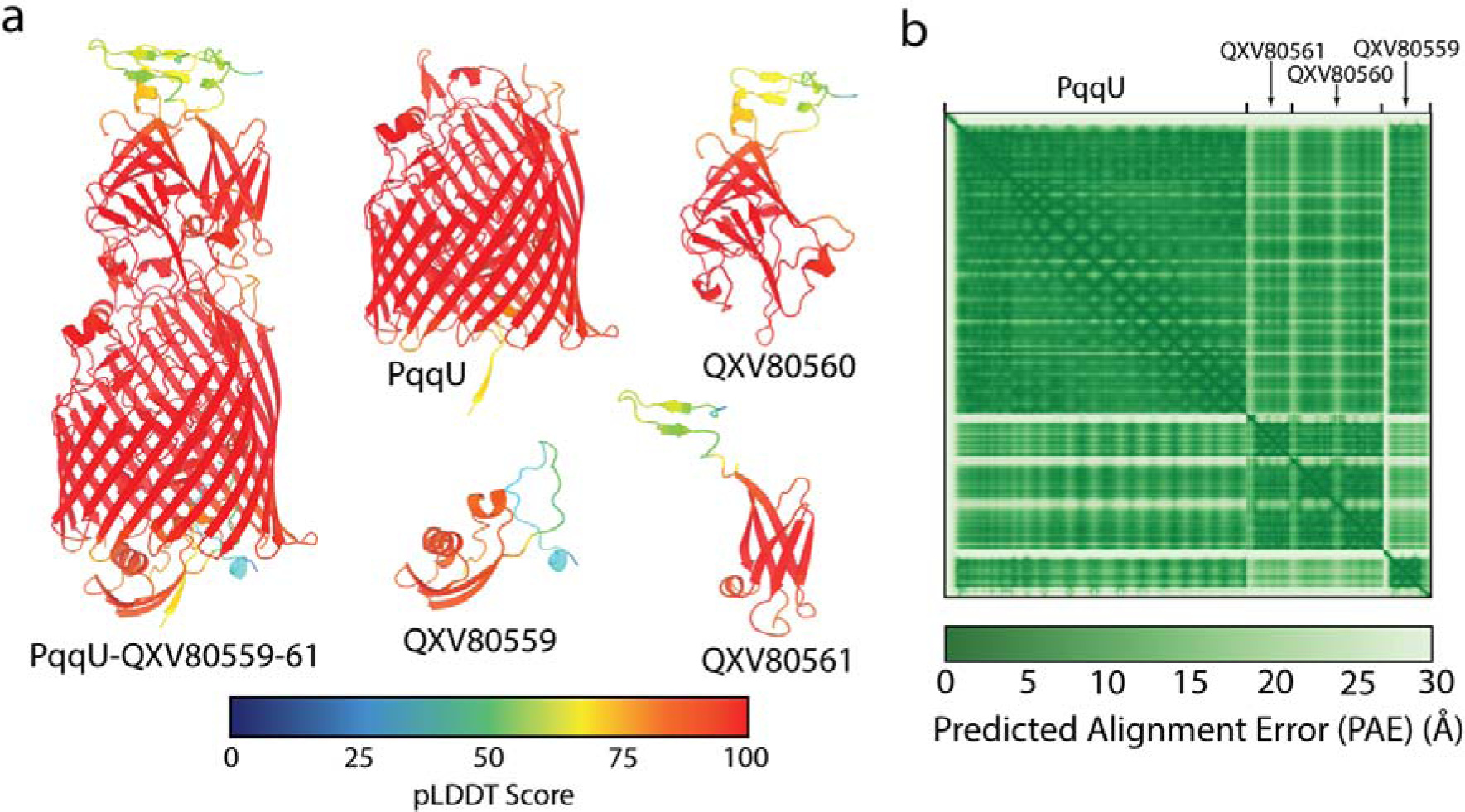
Confidence scores for the AlphaFold2 model of the PqqU-IsaakIselin Phage receptor binding protein complex. (a) The PqqU and phage binding proteins (QXV80560-62) are coloured by pLDDT confidence score. (b) 2D matrix of predicted alignment error (PAE) for the PqqU-QXV80560-62 structure. ***See attached file (too large to embed in text)***

**Figure S6:**
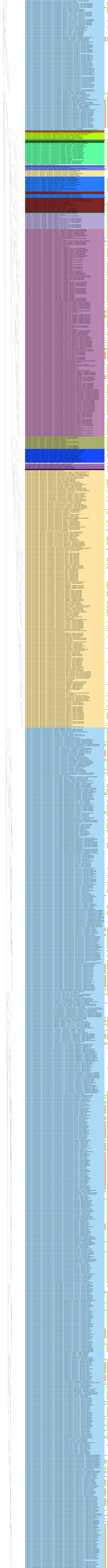
Horizontal genome tree of PqqU producers. Phylogenetic tree constructed from a concatenated alignment of 16 ribosomal proteins. The tree includes 1,861 genomes where 8 or more ribosomal proteins were identified. Genome names are based on their GTDB-Tk classification found in.

**Figure S7:**
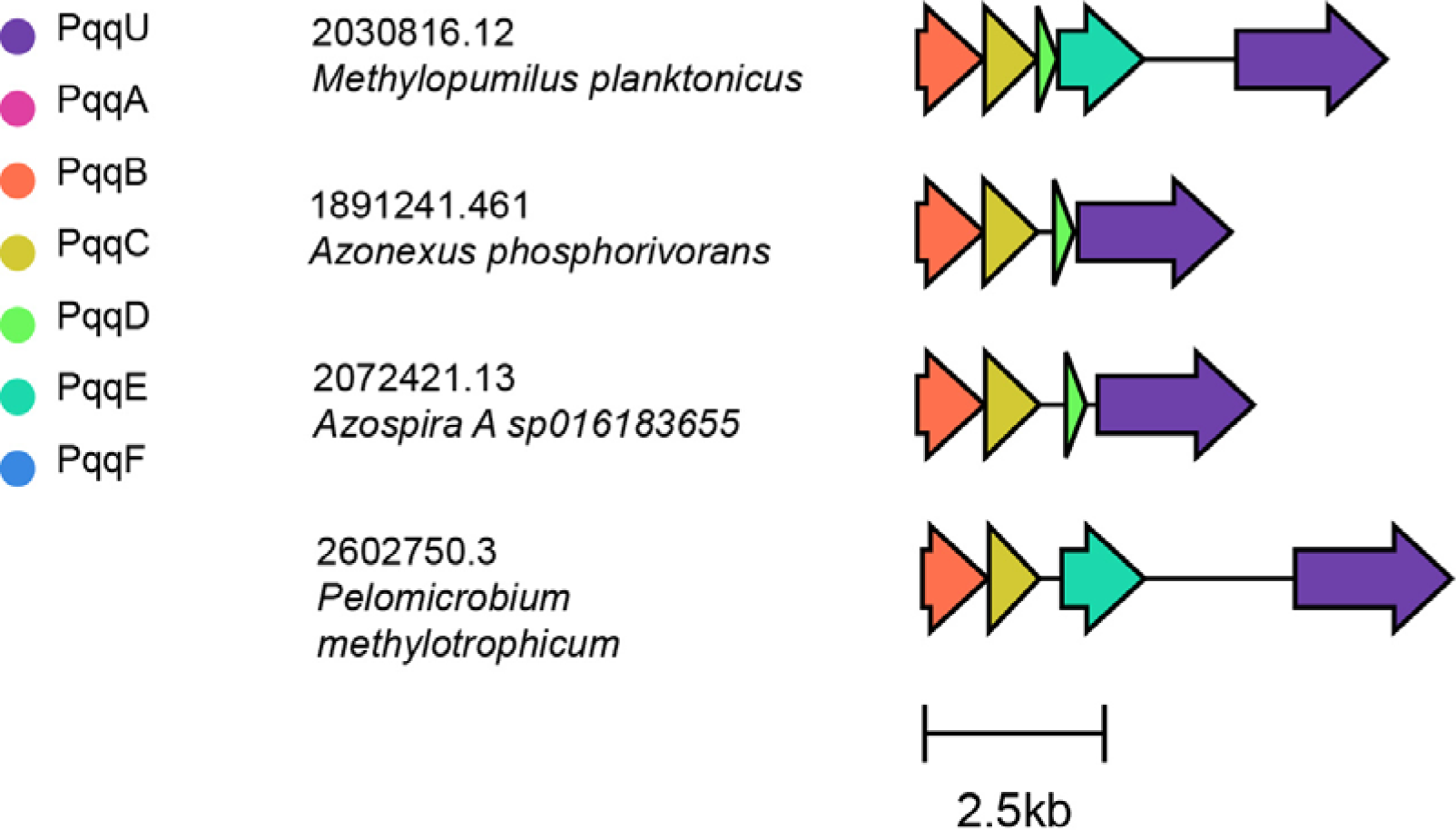
Synteny and orientation of PqqU and the PQQ biosynthesis operon in four *Burkholderiales* genomes.

## SUPPLEMENTAL TABLES

**Table S1:**
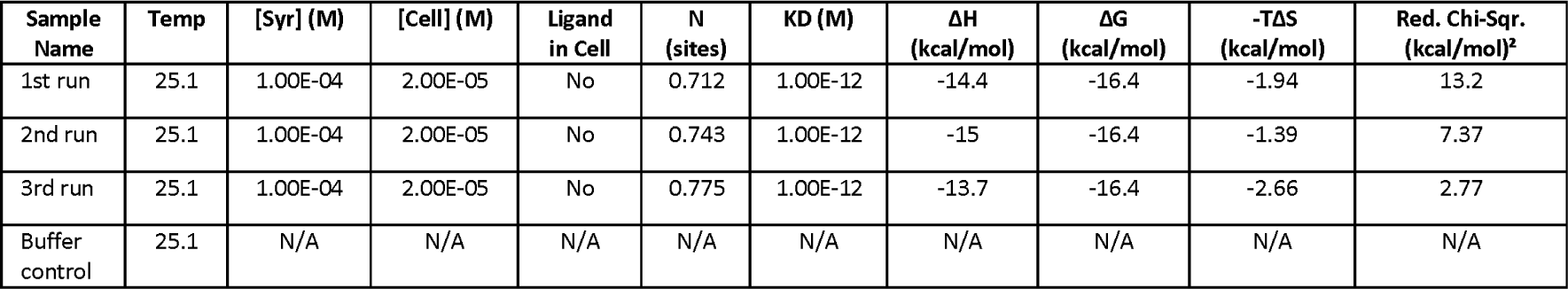
ITC-derived PqqU-PQQ binding parameters. ITC experiment output for three replicate runs and the buffer control. Output generated by the MicroCal Analysis software.

**Table S2:**
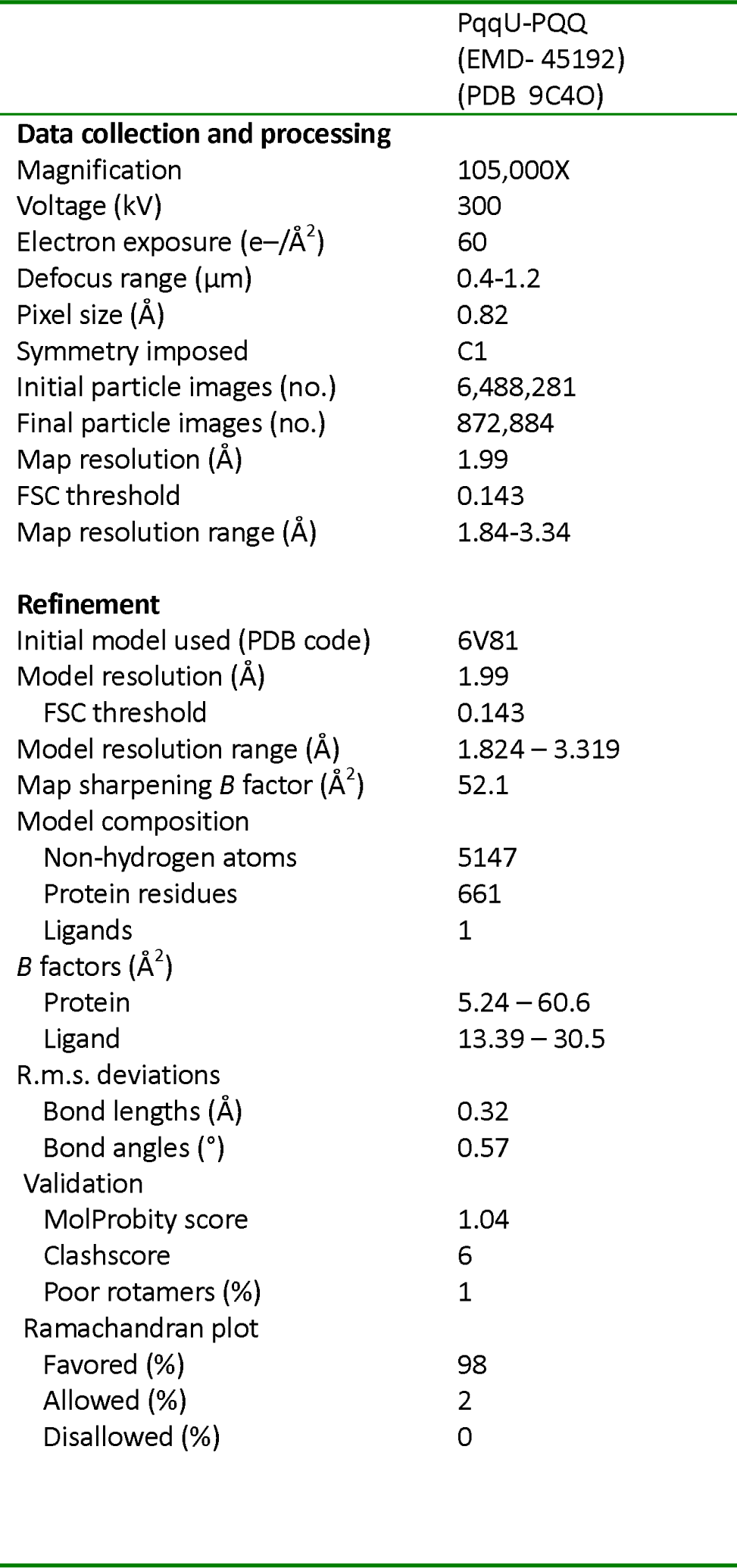
Cryo-EM data collection, refinement and validation statistics. All datasets were collected with a zero-loss filtering slit width of 10 eV and with 60 frames per movie.

**Table S3:**
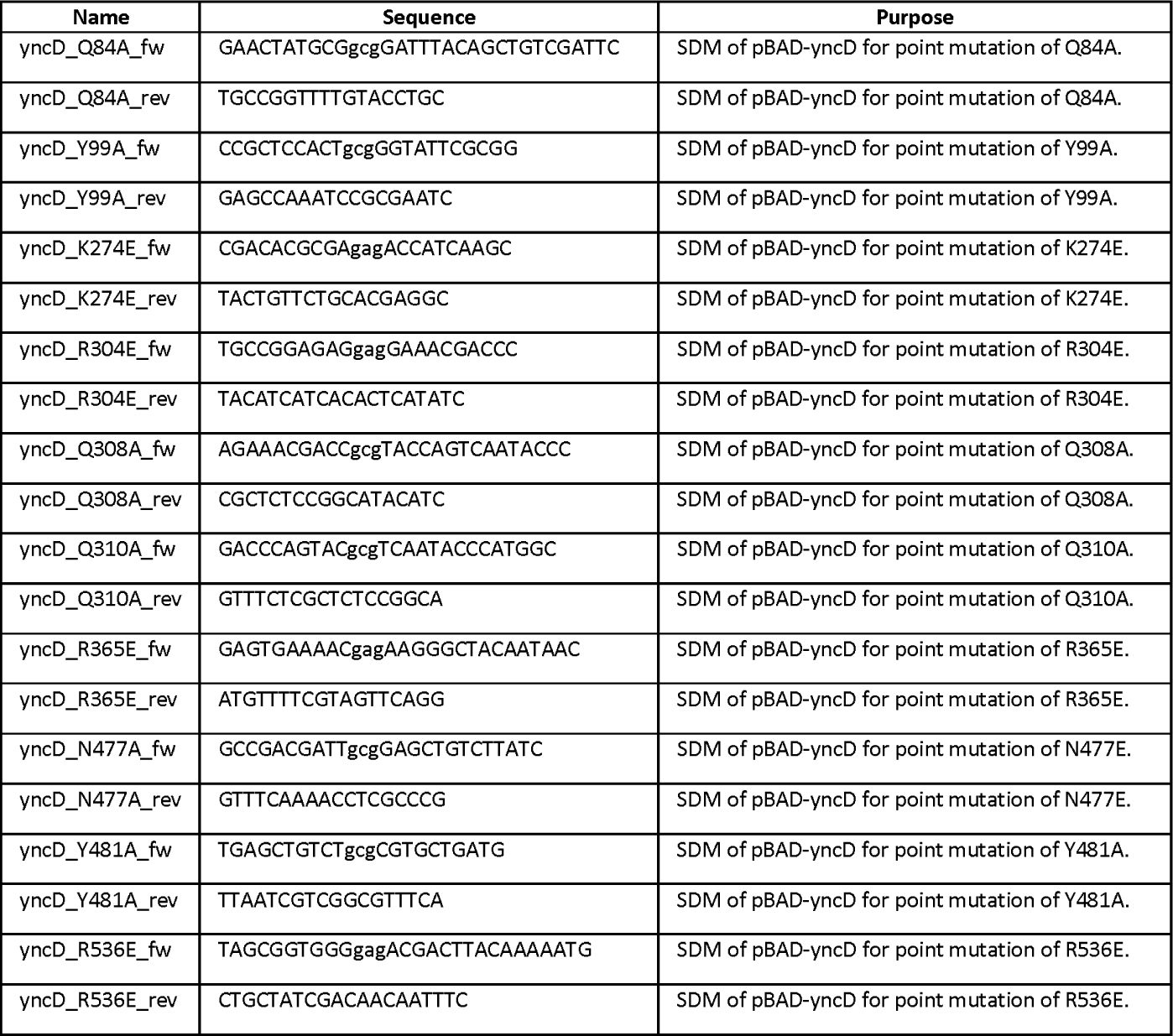
Primer table. Primers used in this study to perform site-directed mutagenesis (SDM). Reverse primers (designated ‘rev’) are shown as reverse complement sequences.

**Table S4:**
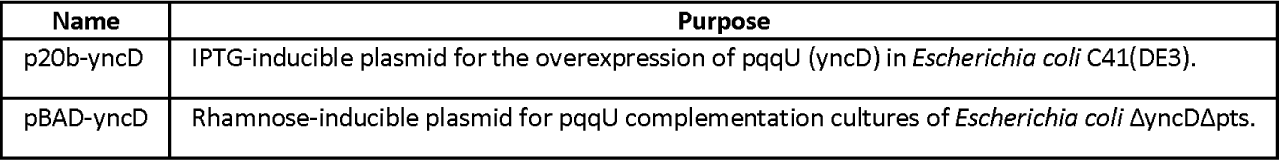
Plasmid table. Plasmids used in this study.

**Table S5:**
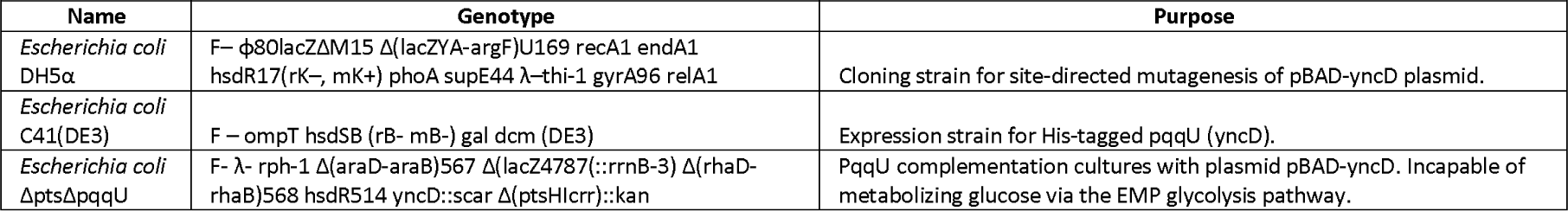
Strain table. Bacterial strains used in this study.

## SUPPLEMENTAL MOVIES

**Movie S1: Contouring of coulomb potential map for the PqqU-PQQ structure**

**Movie S2: Morph of conformational changes in loops 7 and 8 between apo and PQQ bound PqqU**

**Movie S3: Morph of conformational changes in the PqqU binding pocket sidechains between apo and PQQ bound PqqU**

## SUPPLEMENTAL DATA

**Supplemental data S1: AF2 models of the PqqU-phage proteins complex**

**Supplemental data S2: Full output sheet of PqqU homologues associated metadata and quinoproteins identified**

**Supplemental data S3: Multiple alignment file of PqqU homologs**

